# BRIDGE: A broad-host platform for interkingdom DNA delivery enabling the genetic domestication of phylogenetically diverse yeasts

**DOI:** 10.64898/2026.09.17.752325

**Authors:** Reem Swidah, Ryan R. Cochrane, Fernando Valle, Daniela Delneri

**Affiliations:** Manchester Institute of Biotechnology, The University of Manchester, 131 Princess Street, Manchester M1 7DN, UK; Division of Evolution, Infection, and Genomics, Michael Smith Building, University of Manchester, Dover Street, Manchester, M13 9PT, UK; BP Biosciences Centre, San Diego, CA 92121, USA

## Abstract

Efficient DNA delivery remains one of the greatest barriers to engineering non-conventional yeasts, limiting their genetic domestication and exploitation as next-generation microbial cell factories. Here, we present **BRIDGE** (Bacteria-to-yeast Rapid Interkingdom DNA Gene Exchange), an integrated synthetic biology platform for broad-host interkingdom DNA delivery. BRIDGE is centred on a compact 6-kb Pan/ARS–oriT broad-host BRIDGE vector, enabled by a super conjugative helper plasmid (pSC5). Using BRIDGE, we expanded interkingdom DNA transfer from representatives of four genera accessible using pSC5 alone to ten phylogenetically diverse yeast genera, including *Saccharomyces, Maudiozyma, Starmerella, Kazachstania, Yarrowia, Zygosaccharomyces, Lachancea, Kluyveromyces, Pichia* and *Komagataella*. To demonstrate the versatility of BRIDGE, we engineered a 16-kb visual reporter carrying a synthetic five-gene violacein biosynthetic pathway from *Chromobacterium violaceum*. Individual transcriptional units were first constructed using the YeastFab modular cloning system and subsequently assembled into the complete pathway by single-step *in vivo* homologous recombination in *Saccharomyces cerevisiae*, followed by rapid plasmid recovery using the EASY-C platform. The reporter was successfully delivered to all ten yeast genera, demonstrating efficient BRIDGE transfer of complete multigene synthetic pathways. Robust violacein production was observed across all six *Saccharomyces* species tested, providing a rapid visual marker for transformant identification, while modest functional pathway expression was also detected in *Kazachstania* and *Kluyveromyces*. Successful pathway transfer, albeit without visible pigmentation, was achieved in the remaining genera indicating that regulatory compatibility, rather than DNA transfer, is the principal determinant of heterologous pathway expression in distantly related yeasts. Collectively, BRIDGE establishes an integrated Design–Build–Recover–Deliver workflow that combines modular *in vitro* construction of transcriptional units, single-step *in vivo* assembly of multigene pathways, rapid plasmid recovery using the EASY-C platform and broad-host interkingdom DNA delivery. This versatile framework enables the rapid genetic modification of previously intractable microorganisms, expanding the synthetic biology toolbox for sustainable biomanufacturing and industrial biotechnology.

Abstract Figure.
Overview of the BRIDGE technology
BRIDGE (Bacteria-to-yeast Rapid Interkingdom DNA Gene Exchange) is a broad-host platform for conjugation-mediated DNA transfer from bacteria to phylogenetically diverse yeasts. An *Escherichia coli* donor cell (left) harbours a ~60 kb super conjugative helper plasmid encoding the conjugative machinery together with a Pan/ARS-based BRIDGE delivery vector capable of stable replication across diverse yeast species. The vector carries an oriT sequence that mediates interkingdom DNA transfer and a five-gene violacein (*vioA–E*) biosynthetic pathway that serves as a visual reporter of successful DNA delivery. Following transfer, recipient yeast cells (right) acquire the delivery vector and develop varying intensities of purple pigmentation, enabling rapid identification of positive colonies. BRIDGE was successfully evaluated across 12 phylogenetically diverse yeast species representing 10 genera, including *Saccharomyces cerevisiae, Maudiozyma bulderi* 39, *Starmerella batistae, Starmerella bombicola, Kazachstania naganishii, Kazachstania aerobia, Yarrowia lipolytica, Zygosaccharomyces bailii, Lachancea thermotolerans, Kluyveromyces aestuarii, Pichia manshurica*, and *Komagataella phaffii*.

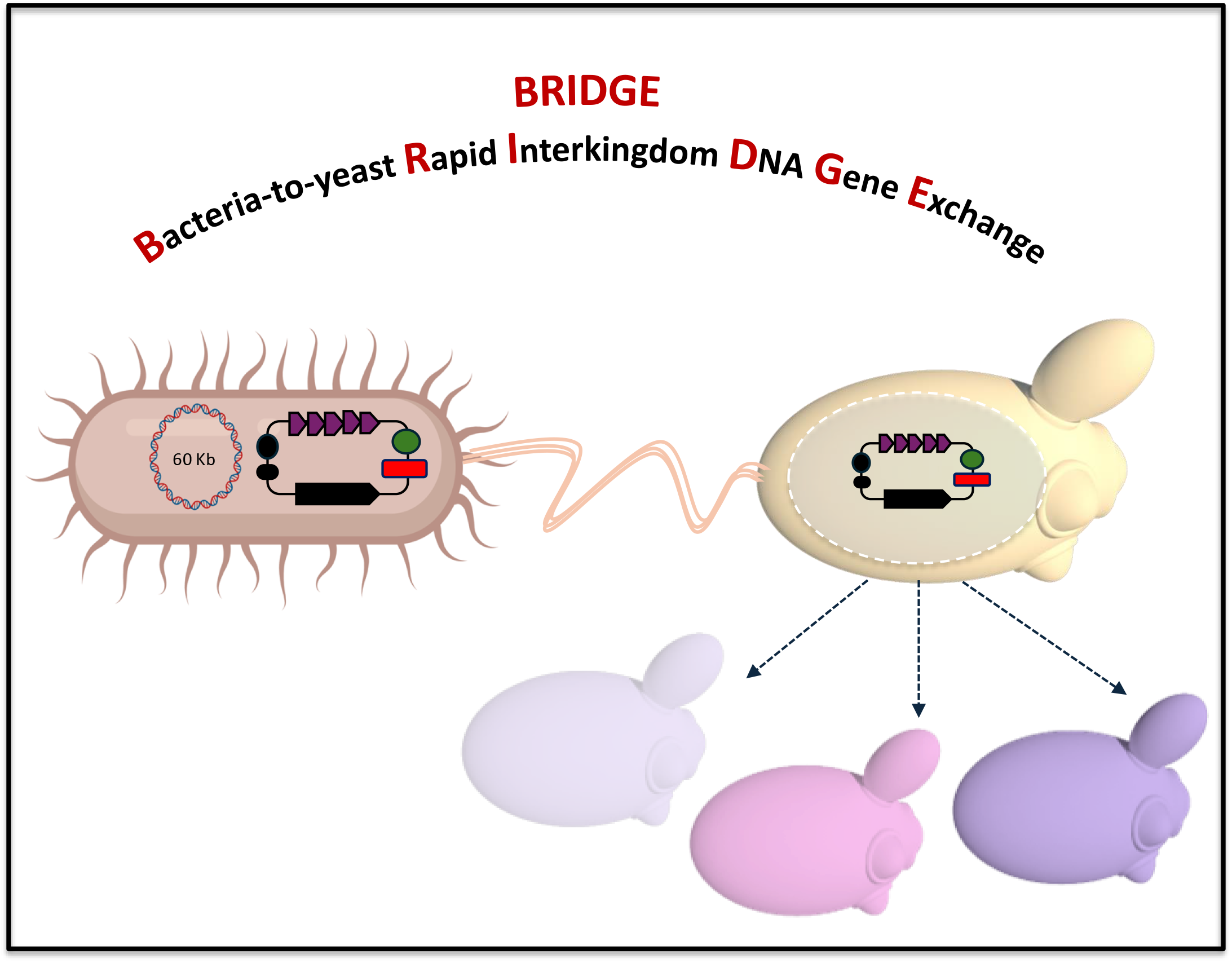

## 1. Introduction

Yeasts have become indispensable platforms for synthetic biology, metabolic engineering and sustainable biomanufacturing. While *Saccharomyces cerevisiae* remains the principal model organism, an increasing number of non-conventional yeasts are emerging as next-generation microbial cell factories owing to their unique physiological and metabolic capabilities. Species such as *Starmerella bombicola, Starmerella batistae, Kazachstania naganishii, Kazachstania aerobia* and *Maudiozyma bulderi* exhibit desirable industrial traits, including growth under low-pH conditions, tolerance to environmental stresses, and the production of high-value compounds such as organic acids, acetate esters, sophorolipids and other specialty chemicals (Alfian, Watchaputi et al. 2022, Balarezo-Cisneros, Timouma et al. 2023, Lin, Walker et al. 2023, Bigotto, Dahmer et al. 2025). Despite their considerable potential, many non-conventional yeasts remain genetically intractable because efficient and broadly applicable genetic engineering tools are still lacking (Wagner and Alper 2016, Lobs, Schwartz et al. 2017).

A major contributor to this genetic intractability is inefficient DNA delivery, which is a prerequisite for robust genome engineering, metabolic engineering and synthetic biology. Conventional transformation methods, including lithium acetate transformation, electroporation and protoplast transformation, frequently require extensive species-specific optimisation and often perform poorly in non-conventional yeasts (Wagner and Alper 2016, Lobs, Schwartz et al. 2017). Consequently, introducing synthetic DNA into many emerging yeast species remains a major bottleneck that limits their genetic domestication, biological investigation, and wider exploitation for biotechnology.

Bacterial conjugation offers an attractive alternative to conventional transformation by enabling active DNA transfer through the bacterial Type IV secretion system (T4SS) without requiring recipient competence (Smillie, Garcillan-Barcia et al. 2010, Christie, Whitaker et al. 2014). The modern era of interkingdom conjugation-mediated DNA transfer began in the early 1980s with the development of *Agrobacterium tumefaciens*-mediated transformation, which exploits a specialised T4SS to deliver transfer DNA (T-DNA) randomly into plant genomes. This technology established the first practical example of bacterial DNA transfer across kingdom boundaries and revolutionised plant genetic engineering (Gelvin 2003). Inspired by this breakthrough, Heinemann and Sprague demonstrated in 1989 that conjugative plasmids from *Escherichia coli* could directly transfer DNA into *S. cerevisiae*, providing the first evidence that bacterial conjugation could be harnessed for interkingdom DNA delivery to yeast (Heinemann and Sprague 1989). The approach was subsequently extended to additional fungal species, and more recently, engineered super conjugative plasmids expanded DNA transfer to selected fungi, including *S. cerevisiae, Metschnikowia gruessii*, and the emerging pathogen *Candida auris* (Cochrane, Shrestha et al. 2022). Despite these advances, the host range has remained relatively narrow, and broadly applicable expression vectors capable of stable replication and maintenance across phylogenetically diverse yeasts have not been available.

Here, we present BRIDGE (Bacteria-to-Yeast Rapid Interkingdom DNA Gene Exchange), an integrated synthetic biology platform for broad-host interkingdom DNA delivery across phylogenetically diverse yeasts. Building on the previously developed super-conjugative helper plasmid pSC5, we sought to determine whether separating the conjugation machinery from the transferable genetic cargo could provide a more flexible and efficient approach to DNA delivery. Rather than transferring and maintaining the large helper plasmid containing conjugation machinery that is not required in the yeast recipient, BRIDGE delivers the genetic cargo on a smaller, independent expression vector. This separation could improve DNA transfer while facilitating the construction, recovery and exchange of complex synthetic cargo across diverse yeast hosts.

To implement this strategy, we engineered a compact ~6-kb Pan/ARS–oriT broad-host expression vector, hereafter referred to as the BRIDGE vector, designed to support autonomous plasmid maintenance and synthetic DNA expression across diverse yeast species. Using this vector, we demonstrated interkingdom DNA delivery across ten phylogenetically diverse yeast genera. To evaluate the delivery and functional expression of complex multigene cargo, we further constructed a 16-kb synthetic violacein reporter carrying the complete five-gene biosynthetic pathway from *Chromobacterium violaceum* (August, Grossman et al. 2000, Balibar and Walsh 2006). Together, BRIDGE establishes a unified Design–Build–Recover–Deliver workflow for constructing, recovering and transferring synthetic DNA across diverse yeast hosts, providing a versatile foundation for engineering previously challenging microorganisms for synthetic biology, sustainable biomanufacturing and industrial biotechnology (Fig. 1).

**Figure 1.**
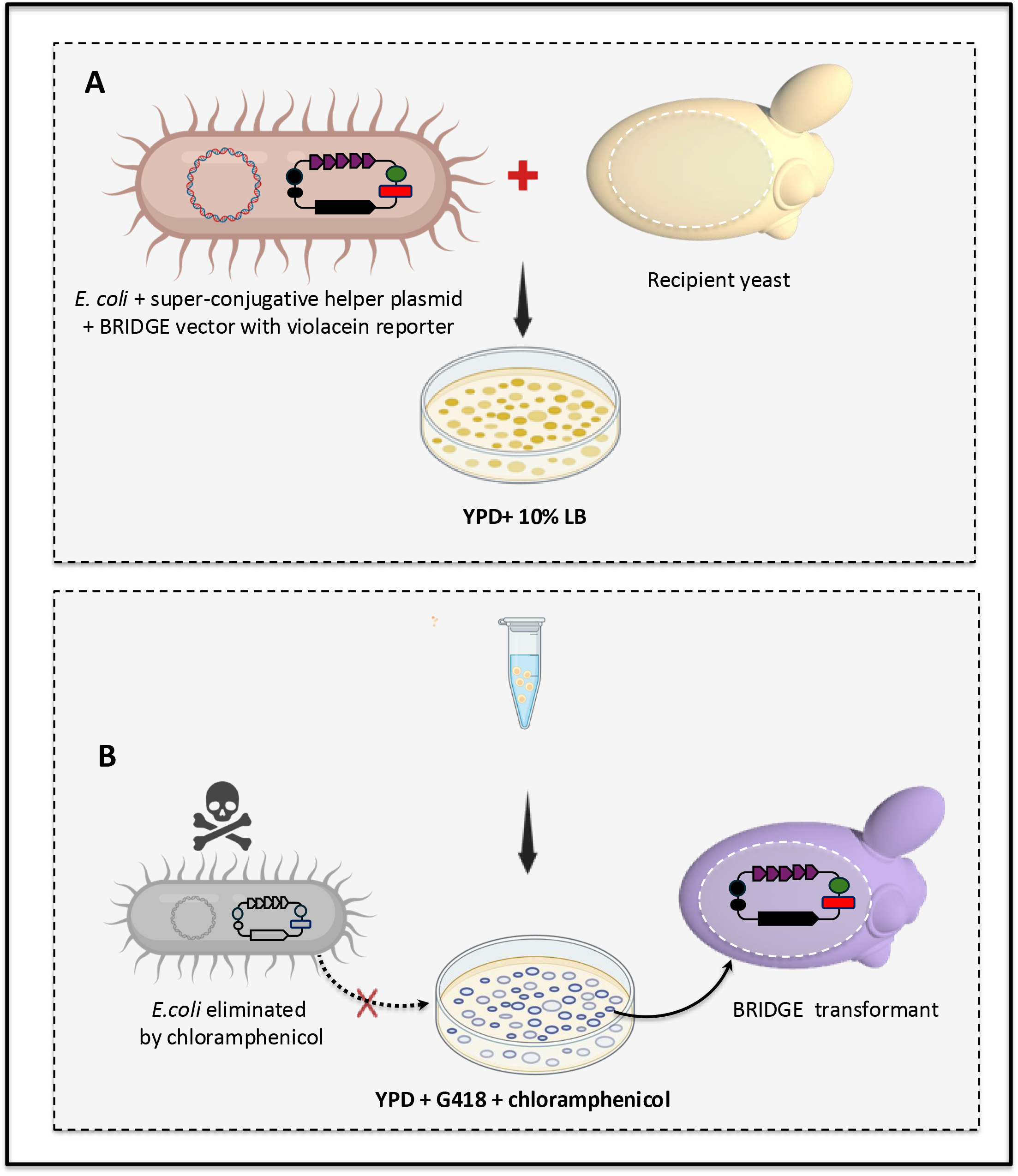
Two-step workflow of the BRIDGE technology. **(A)** Step 1: Interkingdom DNA transfer. An *Escherichia coli* donor strain carrying a super conjugative helper plasmid, pSC5, encoding the conjugation machinery together with a Pan/ARS-based broad-host delivery vector carrying the violacein (*vioA–E*) reporter is mixed with recipient yeast cells. The mixed culture is incubated overnight at 30°C on YPD supplemented with 10% LB, allowing both yeast and bacteria to grow. (**B**) Step 2: Selection and confirmation of yeast transformants. Cells are collected and plated onto YPD supplemented with G418 and chloramphenicol. G418 selects yeast cells that have received the expression vector, while chloramphenicol eliminates the remaining *E. coli* donor cells. Plates are incubated at 30°C, and successful DNA transfer events are first identified by the appearance of purple yeast transformants produced by the violacein reporter (primary screening). Positive transformants are then confirmed by colony PCR to verify successful transfer of the delivery vector (secondary screening).

## 2. Materials and methods

### 2.1 Strains, plasmids, synthetic DNA fragments, and growth media

The yeast strains used in this study included *S. cerevisiae* BY4741, *M. bulderi* 39, *S. batistae, S. bombicola, K. naganishii, K. aerobia, Y. lipolytica, Z. bailii, L. thermotolerans, K. aestuarii, P. manshurica*, and *K. phaffii*, all obtained from the laboratory of Daniela Delneri (Manchester Institute of Biotechnology, University of Manchester, UK). YeastFab plasmids (pHC_Kan_-P, pHC_Kan_-O, pHC_Kan_-T, and pPOT-*LEU2*) were obtained from the laboratory of Yizhi Cai (Manchester Institute of Biotechnology, University of Manchester, UK) (Guo et al., 2015). The origin of replication Pan/ARS of the BRIDGE delivery vector was derived from pUDBzB-41 (Jayaprakash, Barroso et al. 2023). The kanMX (G418 resistance) cassette was obtained from the laboratory of Mark Ashe (Faculty of Biology, Medicine and Health, University of Manchester, UK)(Swidah, Wang et al. 2015). The ~60 kb super conjugative helper plasmid, pSC5, was a gift from Bogumil Karas (Addgene plasmid # 188602; http://n2t.net/addgene:188602; RRID:Addgene_188602) and was kindly provided by Dr Ryan R. Cochrane (Faculty of Biology, Medicine and Health, University of Manchester, UK) (Cochrane, Shrestha et al. 2022). Codon-optimized *vioA, vioB, vioE, vioD*, and *vioC* genes were synthesized by Twist Bioscience (USA) and supplied in standard cloning vectors carrying the kanamycin selectable marker. Yeast strains were routinely cultured in YPD medium (10 g L^−1^ yeast extract, 20 g L^−1^ peptone, and 20 g L^−1^ glucose). Selective growth was performed on synthetic complete dextrose (SCD) medium supplemented with 2% glucose and the appropriate selection marker, including G418 (Geneticin; Thermo Fisher Scientific) or amino acid dropout media where required. For experiments involving the violacein reporter, L-tryptophan was supplemented at threefold the standard concentration in minimal medium to enhance pigment production. *E. coli* strains were cultured in Luria–Bertani (LB) medium and selected with carbenicillin (100 μg mL^−1^), kanamycin (50 μg mL^−1^), or gentamicin (100 μg mL^−1^), as appropriate. Unless otherwise stated, media components were obtained from Formedium (UK) or Fisher Scientific (UK).

### 2.2 Bacterial transformation

Plasmids were transformed into chemically competent *E. coli* DH5α cells by heat shock following standard procedures. For plasmids containing the ccdB counter-selection cassette, chemically competent ccdB-resistant *E. coli* cells were used instead. Briefly, competent cells were incubated with plasmid DNA on ice for 30 min, heat shocked at 42°C for 45 s, and recovered in pre-warmed S.O.C. medium (Invitrogen, Thermo Fisher Scientific, Waltham, MA, USA; Cat. No. 15544034) for 1 h at 37°C with shaking (225 rpm). Cells were concentrated by centrifugation and plated onto LB agar supplemented with the appropriate selective antibiotic, followed by overnight incubation at 37°C. Transformation efficiency was monitored using pUC19 as a positive control. For plasmids smaller than 10 kb, DH5α Mix & Go! Competent Cells (Zymo Research, Orange, CA, USA) were used according to the manufacturer’s instructions, except that the competent cells were prepared at approximately threefold higher concentration than recommended to improve transformation efficiency. When transforming plasmids containing the ccdB cassette, ccdB Survival Competent Cells (Zymo Research) were used according to the manufacturer’s instructions.

### 2.3 Yeast transformation and construct confirmation

The complete BRIDGE delivery vector was assembled *in vivo* in *S. cerevisiae* by lithium acetate/polyethylene glycol (LiAc/PEG)-mediated transformation as previously described (Gietz and Woods 2002). Transformants were selected on YPD agar supplemented with G418 (400 μg mL^−1^). Five to eight independent transformants were screened by colony PCR using four junction-specific primer pairs spanning adjacent assembly junctions to confirm correct vector assembly. Transformants from each yeast species were plated onto YPD agar supplemented with the species-specific concentration of nourseothricin for selection of the pSC5 plasmid or the species-specific concentration of G418 for selection of the BRIDGE broad-host expression vector. The optimal selective concentration of each antibiotic for each yeast species was determined before transformation by spot assays on YPD agar containing a range of antibiotic concentrations (Table S1).

### 2.4 Construction of a YeastFab-compatible BRIDGE delivery vector

A compact YeastFab-compatible broad-host destination vector (~6 kb) was engineered to serve as the transferable DNA cargo for the BRIDGE platform. The vector was designed to support functionalities as a multi-host expression shuttle vector, including *E. coli*, conventional and non-conventional yeasts, and modular assembly of synthetic genetic cargo for interkingdom DNA transfer. The plasmid backbone comprised a pan-fungal autonomously replicating sequence (Pan/ARS) to enable plasmid maintenance in *S. cerevisiae* and diverse non-conventional yeast species (Jayaprakash, Barroso et al. 2023), an origin of transfer (oriT) to permit mobilisation by the pSC5 plasmid (Cochrane, Shrestha et al. 2022), a kanMX cassette conferring G418 resistance for yeast selection (Swidah, Wang et al. 2015), and an AmpR/ColE1 module for plasmid propagation and selection in *E. coli*. To enable modular cloning, the vector incorporated a *ccdB* counter-selection cassette flanked by Type IIS restriction enzyme recognition sites, allowing subsequent replacement of the cassette with standardized YeastFab promoter, open reading frame (ORF), and terminator modules through Golden Gate assembly (Guo, Dong et al. 2015, Garcia-Ruiz, Auxillos et al. 2018). This design allows rapid construction of either single transcriptional units or complex multigene pathways using standardised YeastFab DNA parts. All DNA fragments were assembled directly *in vivo* in *S. cerevisiae* through homologous recombination using approximately 60-bp homologous sequences at both termini to direct recombination with adjacent fragments, enabling seamless assembly of the complete plasmid backbone. Successful *in vivo* assembly was evaluated using four junction PCR assays spanning adjacent assembly junctions. Following *in vivo* assembly, plasmids were recovered from yeast using the previously described EASY-C (Extraction and Analysis of Small Yeast Chromosomes) workflow (Swidah, Monti et al. 2026). Correctly assembled plasmids were recovered using EASY-C and sequence verified as described in Sections 2.8–2.9. Only sequence-verified plasmids were used for all subsequent experiments. The resulting sequence-verified plasmid served as a reusable YeastFab-compatible destination vector for downstream insertion of standardized genetic modules and biosynthetic pathways, providing the transferable DNA cargo for all subsequent BRIDGE-mediated interkingdom DNA transfer experiments.

### 2.5 Construction of a BRIDGE delivery vector carrying the synthetic violacein pathway

The violacein biosynthetic pathway comprising *vioA, vioB, vioE, vioD*, and *vioC* was assembled on the BRIDGE delivery vector using the YeastFab modular cloning system. Individual transcriptional units consisting of a promoter, coding sequence, and terminator were first constructed by Type IIS Golden Gate assembly. Promoters and terminators were selected from *S. cerevisiae, Saccharomyces paradoxus*, and *Saccharomyces eubayanus* to provide constitutive expression of each biosynthetic gene. Coding sequences were codon-optimised before cloning. Regulatory elements were cloned into the pHC_KAN_ vector, whereas coding sequences were cloned into pHC_KAN_-O. Individual transcriptional units were confirmed by colony PCR and Sanger sequencing. The complete violacein pathway was assembled *in vivo* in *S. cerevisiae* by homologous recombination using 60-bp homologous flanking regions between adjacent DNA fragments. The linearized BRIDGE vector backbone and PCR-amplified transcriptional units were co-transformed into yeast, and transformants were selected on YPD agar supplemented with G418. Colonies exhibiting the characteristic purple phenotype after approximately two days were selected as primary candidates for correct pathway assembly. Eight independents purple transformants were analysed by six junction PCR assays, confirming correct assembly of the complete pathway in all isolates. Correctly assembled BRIDGE vectors were recovered directly from yeast using the EASY-C workflow as described in Sections 2.7–2.8. Sequence-verified vectors were subsequently used for construction of BRIDGE donor strains and all downstream BRIDGE-mediated horizontal DNA transfer experiments.

### 2.6 Construction of the *Escherichia coli* donor strain for BRIDGE-mediated DNA transfer

The sequence-verified 6-kb BRIDGE broad-host expression vector, recovered from *S. cerevisiae* using the EASY-C workflow and confirmed by full-plasmid Sanger sequencing, was transformed into chemically competent ccdB-resistant *E. coli* cells. Transformants were selected on LB agar supplemented with carbenicillin (100 μg mL^−1^). A confirmed transformant was subsequently mixed with an *E. coli* donor strain harbouring the ~60-kb pSC5 plasmid by streaking both strains together onto an LB agar plate and incubating overnight at 37°C to facilitate horizontal transfer of pSC5 into the *E. coli* strain carrying the BRIDGE vector. Bacterial cells were recovered from the plate in sterile water and plated onto selective LB agar supplemented with carbenicillin (100 μg mL^−1^) and gentamicin (40 μg mL^−1^) to isolate clones carrying both the BRIDGE vector and pSC5. The presence of both plasmids was confirmed by plasmid-specific colony PCR using five independent biological isolates. Confirmed donor strains were preserved as glycerol stocks at −80°C and used for subsequent BRIDGE-mediated DNA transfer experiments. This procedure was repeated using the sequence-verified 16-kb BRIDGE delivery vector harbouring the synthetic violacein biosynthetic pathway to generate the corresponding *E. coli* donor strain for BRIDGE-mediated delivery of the violacein pathway.

### 2.7 Recovery of the assembled vector using EASY-C

Correctly assembled plasmids were recovered from yeast using the Extraction and Analysis of Small Yeast Chromosomes (EASY-C) workflow (Swidah, Monti et al. 2026). Briefly, plasmid DNA was isolated from yeast, transformed into chemically competent *E. coli* DH5α cells for propagation, and purified using the QIAprep Spin Miniprep Kit (QIAGEN) according to the manufacturer’s instructions.

### 2.8 Sequence verification

Purified plasmids were verified by full-plasmid Sanger sequencing using the Plasmidsaurus sequencing service. Sequencing reads were aligned to the reference plasmid sequence to confirm complete and accurate plasmid assembly. Only sequence-verified constructs were used for subsequent BRIDGE experiments.

### 2.9 BRIDGE-mediated interkingdom DNA transfer

Donor *E. coli* and recipient yeast strains were prepared as frozen glycerol stocks before BRIDGE experiments. *E. coli* donor strains carrying either the ~60 kb pSC5 plasmid alone, or together with either the empty 6 kb BRIDGE delivery vector or the 16 kb BRIDGE delivery vector harbouring the synthetic violacein reporter were grown overnight in LB medium supplemented with the appropriate antibiotics (gentamicin for strains carrying the pSC5 plasmid alone or gentamicin plus carbenicillin for strains carrying both plasmids). Cultures were diluted to an OD_600_ of 0.01 in fresh selective LB medium and grown to an OD_600_ of approximately 1.0. Cells were harvested by centrifugation (3,000 × *g*, 15 min), resuspended in ice-cold 10% glycerol, aliquoted, and stored at −80°C. Recipient yeast strains were inoculated from single colonies into YPD medium and grown overnight at 30°C. Cultures were diluted to an OD_600_ of 0.2 in fresh YPD containing ampicillin and grown to an OD_600_ of approximately 3.0. Cells were harvested by centrifugation (3,000 × *g*, 5 min), resuspended in ice-cold 10% glycerol, aliquoted, and stored at −80°C. For BRIDGE-mediated DNA transfer, donor and recipient aliquots were thawed on ice and mixed at a ratio of 100 μL donor cells to 50 μL recipient yeast cells. Cell mixtures were spread onto conjugation plates consisting of either 10% LB and rich medium (YPD) or 10% LB and complete minimal glucose medium lacking histidine, both solidified with 2% agar, and incubated at 30°C for 24 h to allow interkingdom DNA transfer. Rich medium (YPD) was used for all non-conventional yeast species, whereas either rich medium or complete minimal glucose medium lacking histidine was used for *S. cerevisiae*. Following co-incubation, cells were recovered from each conjugation plate with 2 mL sterile distilled water, vortexed briefly, and plated onto YPD agar supplemented with chloramphenicol (30 μg mL^−1^) to eliminate residual *E. coli*. Transformants carrying the pSC5 plasmid were selected on YPD containing the species-specific concentration of nourseothricin (NatNT2), whereas transformants carrying the BRIDGE delivery vector were selected on YPD containing the species-specific concentration of G418. The optimal selective concentration of each antibiotic for each yeast species was determined before BRIDGE experiments by spot assays on YPD agar containing a range of antibiotic concentrations. The antibiotic concentrations used for each yeast species are listed in **Table S1**. All BRIDGE experiments were performed using three independent biological replicates for each yeast species. Plates were incubated at 30°C for 3–5 days before transformants were scored. The protocol was adapted from (Cochrane, Shrestha et al. 2022).

### 2.10 Quantification and statistical analysis of BRIDGE transformants

Following BRIDGE-mediated DNA delivery, yeast cells were recovered from mating plates, resuspended in sterile water and plated onto selective YPD agar containing G418 and chloramphenicol. After incubation, colonies were manually counted as described in Section 2.9 to determine the number of BRIDGE transformants. Three independent biological replicates were performed for each yeast species. Plates containing more than 1000 transformants were classified as too numerous to count (TNTC) and recorded as ≥1000 colonies, representing the upper limit of reliable manual enumeration. For countable plates, the mean number of transformants and standard deviation (SD) were calculated from the three biological replicates. Statistical analyses were performed using GraphPad Prism version 10 (GraphPad Software, Boston, MA, USA). Differences in BRIDGE transformant numbers were analysed by one-way analysis of variance (ANOVA) followed by Dunnett’s multiple-comparison test, using *S. cerevisiae* as the reference group. Statistical significance was defined as *P* < 0.05 and is indicated as follows: ns, not significant; *, *P* < 0.05; **, *P* < 0.01; ***, *P* < 0.001; ****, *P* < 0.0001. Species yielding too numerous to count (TNTC; >1000 colonies) were excluded from statistical analyses because accurate colony enumeration was not possible. Species for which no transformants were recovered (0 colonies in all three biological replicates) were also excluded from statistical analyses.

### 2.11 Spot assay

Spot assays were performed to assess violacein production following BRIDGE-mediated delivery of the synthetic reporter pathway. Overnight cultures were grown in selective medium, adjusted to an OD_600_ of 0.10, and subjected to 10-fold serial dilutions. Five microlitres of each dilution were spotted onto YPD agar or YPD agar supplemented with G418 to maintain selection for the BRIDGE delivery vector carrying the synthetic ***vioA–E*** violacein pathway. To evaluate reporter activity under defined growth conditions, cells were also spotted onto synthetic defined (SD) medium supplemented with G418 and a three-fold higher concentration of L-tryptophan to increase the availability of the precursor for violacein biosynthesis. Monosodium glutamate (1 g L^−1^) was used as the nitrogen source in place of ammonium sulfate. Plates were incubated at 30°C for 3–4 days before imaging and phenotypic analysis.

## 3. Results

### 3.1 A diverse yeast panel reveals host-dependent barriers to pSC5-mediated DNA delivery

A recently developed ~60-kb super conjugative helper plasmid pSC5, enables interkingdom DNA transfer from *E. coli* to fungi by carrying the complete conjugative machinery together with selectable markers and a bacterial origin of replication (Cochrane, Shrestha et al. 2022). The plasmid contains *HIS3*, an orthogonally expressed *natNT2* cassette for fungal selection using either the standard or alternative genetic code, and a yeast CEN/ARS sequence for low-copy maintenance. Previous studies demonstrated successful transfer into *S. cerevisiae, M. gruessii, Metschnikowia pulcherrima, Metschnikowia lunata, Metschnikowia borealis, Candida tolerans, Candida bromeliacearum, Candida pseudointermedia, Candida ubatubensis*, and *C. auris* (Cochrane, Shrestha et al. 2022), suggesting that this platform could provide a promising alternative to conventional transformation methods. However, an intact functional plasmid could not be recovered from the fungal recipients, suggesting that stable maintenance or propagation of the plasmid may be limited. We initially sought to determine whether the pSC5 plasmid could overcome the poor transformability of the industrially relevant non-conventional yeast *M. bulderi* CBS8639 (Balarezo-Cisneros, Hanak et al. 2025), which is highly resistant to standard plasmid transformation methods. Although efficient transfer was readily observed in *S. cerevisiae*, no transformants were recovered in *M. bulderi*. We reasoned that this failure could result from the large size of the plasmid, inefficient delivery into recipient cells, or limited plasmid maintenance in this species. To determine whether this represented a broader limitation of the platform, we systematically evaluated its host range across phylogenetically diverse yeasts. Ten yeast genera representing a broad range of conventional and non-conventional species were selected as potential recipients **(Fig. 2A)**. Recipient cells were co-incubated with *E. coli* donor cells carrying the pSC5 plasmid and plated on YPD containing species-specific concentrations of nourseothricin, as described in Section 2.9 (Materials and Methods) **(Fig. 2B)**. Colonies were counted after four days of incubation, and three randomly selected colonies from each positive species were verified by confirmation PCR **(Supplementary Fig. 1)**. DNA transfer was successfully detected in *S. cerevisiae* and species belonging to the genera *Starmerella, Yarrowia*, and *Komagataella*. In contrast, no confirmed transformants were obtained for *M. bulderi, K. naganishii, K. aerobia, Z. bailii, L. thermotolerans, K. aestuarii*, or *P. manshurica*. Across three independent biological replicates, transfer efficiencies varied substantially among the positive species, whereas no transfer events were detected in the remaining recipients **(Fig. 2C)**. These findings demonstrate that, despite its ability to mediate interkingdom DNA transfer, the pSC5 plasmid exhibits a restricted host range and is unable to efficiently access many non-conventional yeasts. We decided to separate the conjugative machinery from the transferable DNA cargo generating a series of conjugative plasmids engineered for improved host accessibility and persistence. Specifically, we reasoned that a substantially smaller plasmid carrying a broad-host replication origin would be transferred and maintained more efficiently than the 60-kb helper plasmid. To test this hypothesis, we engineered a compact BRIDGE delivery vector approximately tenfold smaller than the helper plasmid that retained the origin of transfer (**oriT**) required for mobilisation, while incorporating a pan-fungal autonomously replicating sequence (**PanARS**) to support replication across diverse yeast species. The conjugative machinery remained on the large helper plasmid within the bacterial donor, whereas the newly engineered vector served as the transferable DNA cargo, forming the basis of the BRIDGE platform.

**Figure 2.**
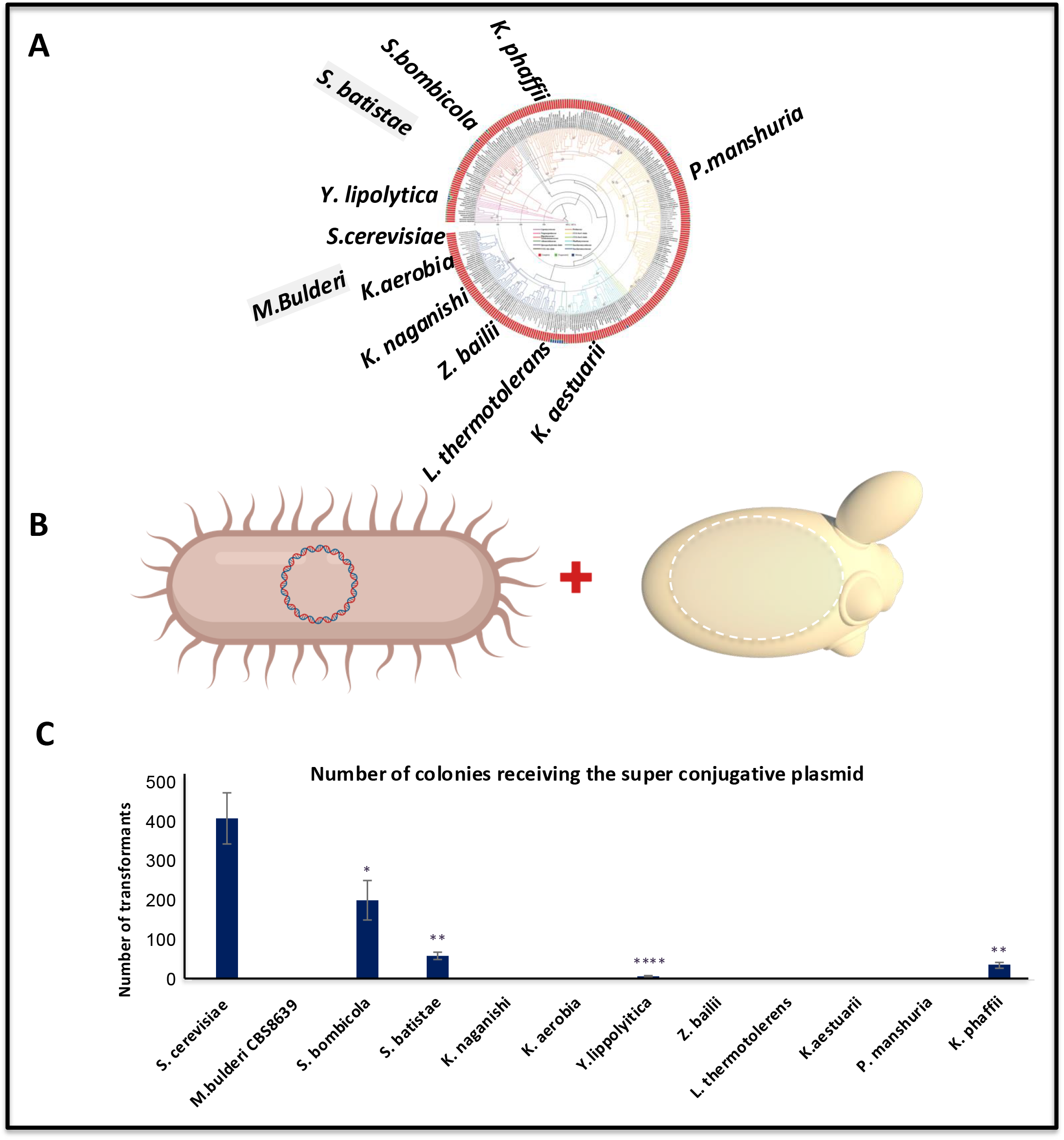
Assessing the efficiency of horizontal transfer of the super conjugative plasmid across 10 yeast genera. **Figure 2. Assessing horizontal transfer of the super conjugative helper plasmid across 10 yeast genera**. **(A)** Phylogenetic tree showing the 12 yeast species representing 10 genera evaluated in this study. The phylogenetic tree was adapted from (Shen, Opulente et al. 2018) (**B**) Schematic of the interkingdom horizontal DNA transfer workflow. An Escherichia coli donor carrying the ~60 kb super conjugative helper plasmid is co-cultured with recipient yeast cells to mediate plasmid transfer. (**C**) Quantification of helper plasmid transfer across 12 yeast species representing 10 genera. Bars represent the mean number of transformants recovered following interkingdom delivery of the super conjugative helper plasmid from three independent biological replicates. Error bars indicate the standard deviation (SD). Statistical significance was determined by one-way ANOVA followed by Dunnett’s multiple-comparison test using *Saccharomyces cerevisiae* as the reference group. Significance is indicated as follows: *, *P* < 0.05; **, *P* < 0.01; ****, *P* < 0.0001. Species for which no transformants were recovered (0 colonies in all three biological replicates) were excluded from statistical analysis.

### 3.2 Engineering a BRIDGE delivery vector

To explore whether separating the conjugation machinery from the transferable DNA cargo could provide a more flexible approach to interkingdom DNA delivery, we engineered a compact ~6-kb broad-host expression/destination vector, hereafter referred to as the BRIDGE vector **(Fig. 3)**. The BRIDGE vector was designed to carry user-defined genetic cargo, support plasmid maintenance across conventional and non-conventional yeasts, and enable conjugative mobilisation by the pSC5 helper plasmid. It was also designed to be compatible with YeastFab, a standardised Type IIS Golden Gate modular cloning framework (Guo, Dong et al. 2015, Garcia-Ruiz, Auxillos et al. 2018), enabling promoter, coding sequence and terminator modules to be assembled into individual transcriptional units and subsequently combined into multigene constructs. The BRIDGE vector contains a pan-fungal autonomously replicating sequence (Pan/ARS) for broad-host plasmid maintenance (Jayaprakash, Barroso et al. 2023), an oriT sequence to enable mobilisation by the pSC5 plasmid (Cochrane, Shrestha et al. 2022), a kanMX cassette for yeast selection (Swidah, Wang et al. 2015), an AmpR/ColE1 module for propagation in *E. coli*, and a ccdB counter-selection cassette that provides the modular cloning entry site for replacement with standardized YeastFab promoter, coding sequence and terminator modules **(Fig. 3A)**. The BRIDGE vector backbone was assembled directly in *S. cerevisiae* by *in vivo* homologous recombination (see Section 2.4). Four junction PCR assays confirmed correct assembly in all five independent transformants tested. The BRIDGE vector was subsequently recovered from yeast using EASY-C **(Fig. 3B)** (Swidah, Monti et al. 2026), propagated in ccdB-resistant *E. coli*, purified and verified by Sanger sequencing (see Section 2.4 for experimental details).

**Figure 3.**
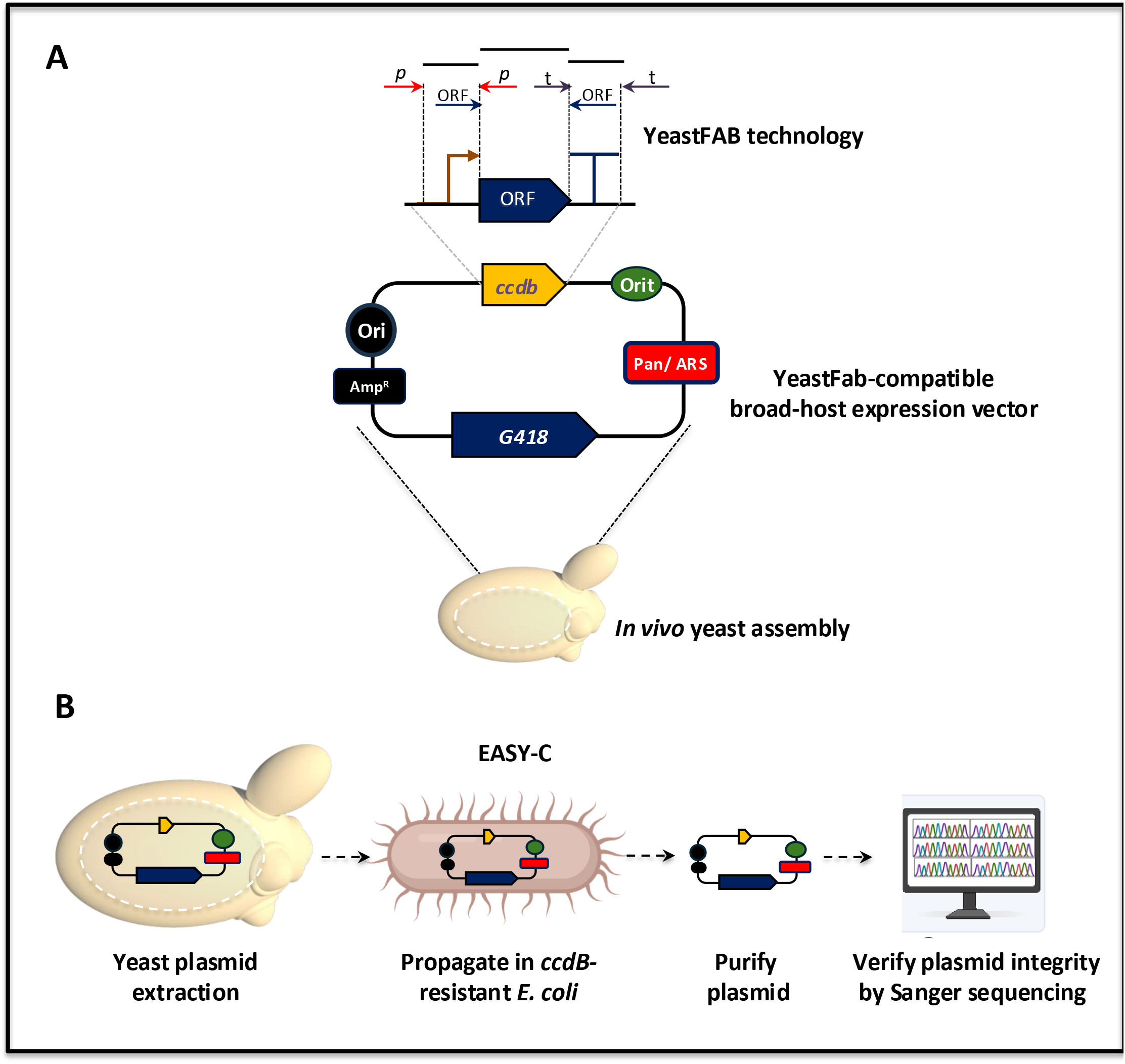
Engineering and validation of a YeastFab-compatible broad-host destination vector for BRIDGE-mediated interkingdom DNA transfer. **(A)** Schematic of the YeastFab-compatible broad-host destination vector. The upper panel illustrates the principle of YeastFab modular cloning, in which standardized promoter (P), open reading frame (ORF), and terminator (T) modules are assembled by one-pot Type IIS Golden Gate cloning through replacement of the ccdB counter-selection cassette (Guo, Dong et al. 2015, Garcia-Ruiz, Auxillos et al. 2018). The resulting destination vector contains a Pan/ARS element for plasmid maintenance in *Saccharomyces cerevisiae* and diverse non-conventional yeasts, an oriT sequence for mobilisation by the super conjugative helper plasmid, a kanMX cassette conferring G418 resistance for yeast selection, and an Amp^R^/ColE1 module for propagation in *Escherichia coli*. The plasmid backbone was assembled *in vivo* in *S. cerevisiae* by homologous recombination. Correct assembly was confirmed by four junction PCR assays performed on five independent biological transformants. **(B)** Validation of the destination vector using the EASY-C workflow. Correctly assembled plasmids were recovered directly from yeast, transformed into *ccdB*-resistant *Escherichia coli*, propagated, purified by plasmid miniprep, and verified by Sanger sequencing before use in subsequent BRIDGE-mediated interkingdom DNA transfer experiments.

### 3.3 BRIDGE expands interkingdom delivery of a BRIDGE delivery vector across diverse yeast genera

To evaluate whether the BRIDGE vector expanded interkingdom DNA transfer, we first constructed an *E. coli* donor strain harbouring both the pSC5 plasmid and the broad-host destination/delivery vector (see Section 2.6). The resulting donor strain was subsequently used for BRIDGE-mediated DNA transfer into representatives of ten yeast genera (see Section 2.9**) (Fig. 4A)**. Following co-cultivation, recipient cells were selected on species-specific G418-containing media and incubated at 30°C for 4– 5 days to recover transformants. While pSC5 successfully transferred cargo to 4 of 10 genera tested **(Fig. 2)**, the **BRIDGE** vector was successfully delivered to representatives of all ten yeast genera examined **(Fig. 4B)**. Successful DNA transfer was observed in *M. bulderi* CBS8639, *S. batistae, S. bombicola, K. naganishii, K. aestuarii, L. thermotolerans, Y. lipolytica, P. manshurica, Z. bailii* and *K. phaffii*, whereas transfer to *K. aerobia* occurred at lower frequency. BRIDGE-mediated DNA transfer was performed using three independent biological replicates for each species, and the mean number of transformants recovered is shown in **Fig. 4C**. To verify successful plasmid delivery, up to five independent transformants recovered from each species were analysed by confirmation PCR. All analysed transformants generated the expected amplification products, confirming successful delivery and maintenance of the BRIDGE vector and demonstrating a high frequency of genuine transformants. During optimisation of the protocol, additional colonies occasionally appeared following prolonged incubation of selective plates at room temperature. Although some represented authentic transformants, prolonged incubation was also associated with the appearance of false-positive colonies of smaller size. Consequently, only colonies recovered within the standard incubation period or confirmed by PCR were included in subsequent analyses. Collectively, these results demonstrate that BRIDGE substantially expands interkingdom DNA delivery by enabling efficient mobilisation of the BRIDGE vector across representatives of ten yeast genera.

**Figure 4.**
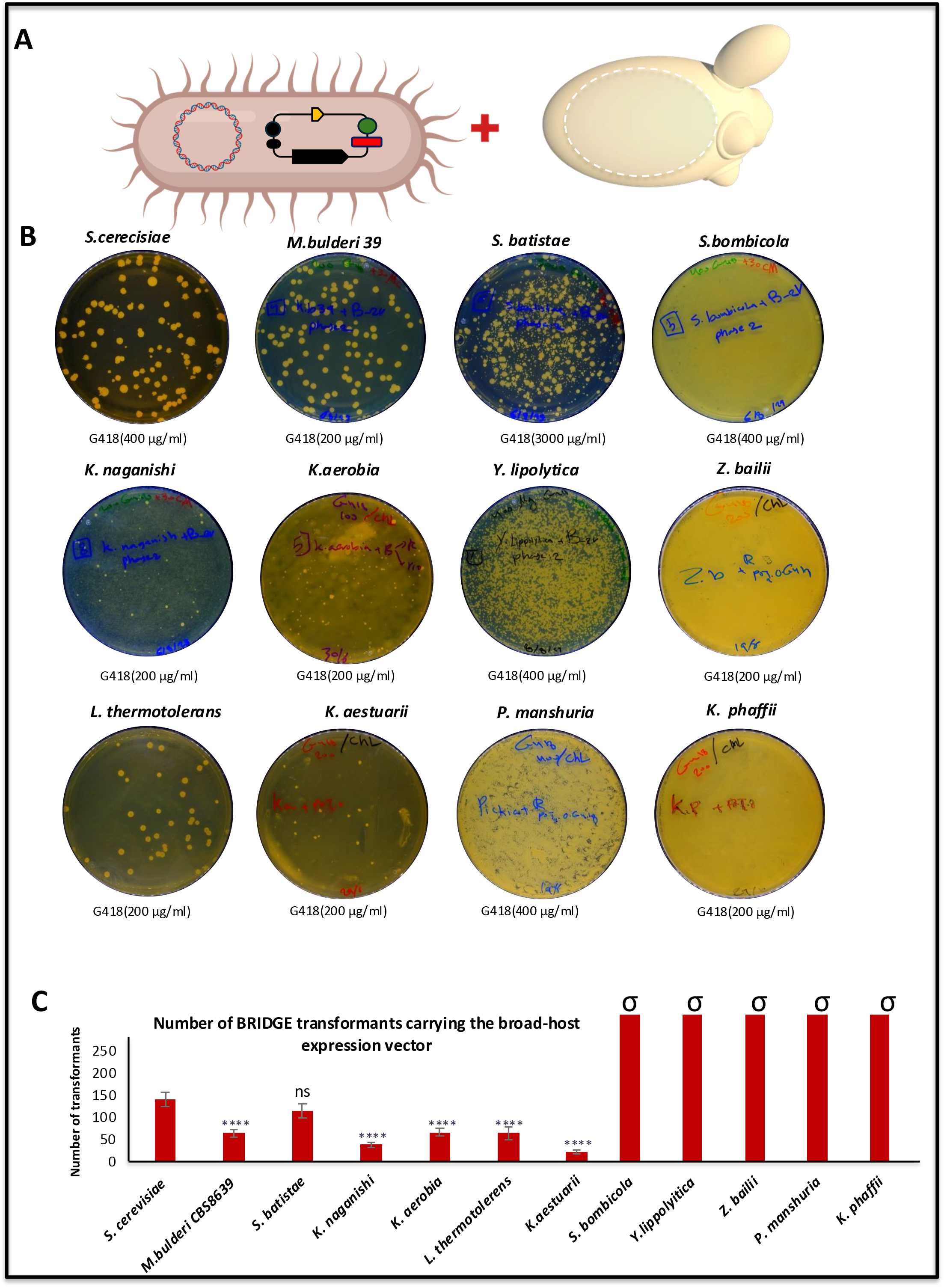
Efficiency of BRIDGE-mediated horizontal transfer of the PanARS expression vector across 10 yeast genera. **Figure 4. BRIDGE expands interkingdom delivery of a BRIDGE delivery vector across representatives of ten yeast genera**. **(A)** Schematic of the BRIDGE interkingdom DNA transfer workflow. An *Escherichia coli* donor carrying the ~60 kb super conjugative helper plasmid together with the 6-kb BRIDGE delivery vector was co-cultured with recipient yeast cells representing 12 species across 10 genera to mediate horizontal DNA transfer. **(B)** Representative transformants obtained four days after BRIDGE-mediated horizontal transfer into *Saccharomyces cerevisiae, Maudiozyma bulderi* 39, *Starmerella batistae, Starmerella bombicola, Kazachstania naganishii, Kazachstania aerobia, Yarrowia lipolytica, Zygosaccharomyces bailii, Lachancea thermotolerans, Kluyveromyces aestuarii, Pichia manshurica*, and *Komagataella phaffii*. Transformants were selected on YPD agar supplemented with chloramphenicol (30 μg mL^−1^) to eliminate *E. coli* and the appropriate concentration of G418 for each yeast species, as indicated beneath each plate. Successful delivery of the BRIDGE vector was confirmed by colony PCR using three to five independent transformants for each species. **(C)** Quantification of BRIDGE-mediated horizontal DNA transfer across 12 yeast species representing 10 genera. Bars represent the mean number of BRIDGE transformants recovered following of the 6-kb BRIDGE delivery vector from three independent biological replicates. Error bars indicate the standard deviation (SD). ****, *P* < 0.0001; ns, not significant. Species yielding >1,000 transformants formed confluent lawns rather than discrete colonies and were therefore excluded from statistical analysis. These samples are indicated by the symbol (σ) above the corresponding bars.

### 3.4 Engineering a violacein-based visual reporter for rapid identification of BRIDGE transformants

To develop a visual reporter and to test whether a large pathway can be transferred between species, we constructed a ~10-kb synthetic cargo containing the complete five-gene violacein biosynthetic pathway (*vioA–E*) from *Chromobacterium violaceum* (August, Grossman et al. 2000, Balibar and Walsh 2006) using the YeastFab modular cloning system **(Fig. 5A,B)**. The *vioA–E* coding sequences were codon-optimised for *S. cerevisiae* (Liu, Luo et al. 2018) and refactored with constitutive regulatory elements derived from *S. cerevisiae, S. paradoxus* and *S. eubayanus* **(Fig. 5C)**. Further details of the vector construction are provided in Materials and Methods Section 2.5. The complete violacein pathway was assembled *in vivo* in *S. cerevisiae* by homologous recombination using 60-bp overlaps between adjacent DNA fragments **(Fig. 5D)**. The linearised BRIDGE vector and PCR-amplified transcriptional units were co-transformed into yeast, and G418-resistant colonies producing the characteristic purple pigment were selected after approximately two days **(Fig. 5E)**. Eight independent purple colonies were analysed using six junction PCR assays, confirming the complete *vioA–E* pathway in all clones. The assembled BRIDGE vectors were then recovered directly from yeast using the EASY-C workflow **(Fig. 5F)**, propagated in *E. coli*, purified and verified by full-plasmid Sanger sequencing. Sequence-verified vectors were subsequently used to generate bacterial donor strains for BRIDGE-mediated DNA transfer experiments. The ~10-kb five-gene pathway therefore provided a pathway-scale cargo for evaluating BRIDGE-mediated DNA delivery and functional expression across diverse yeast hosts.

**Figure 5.**
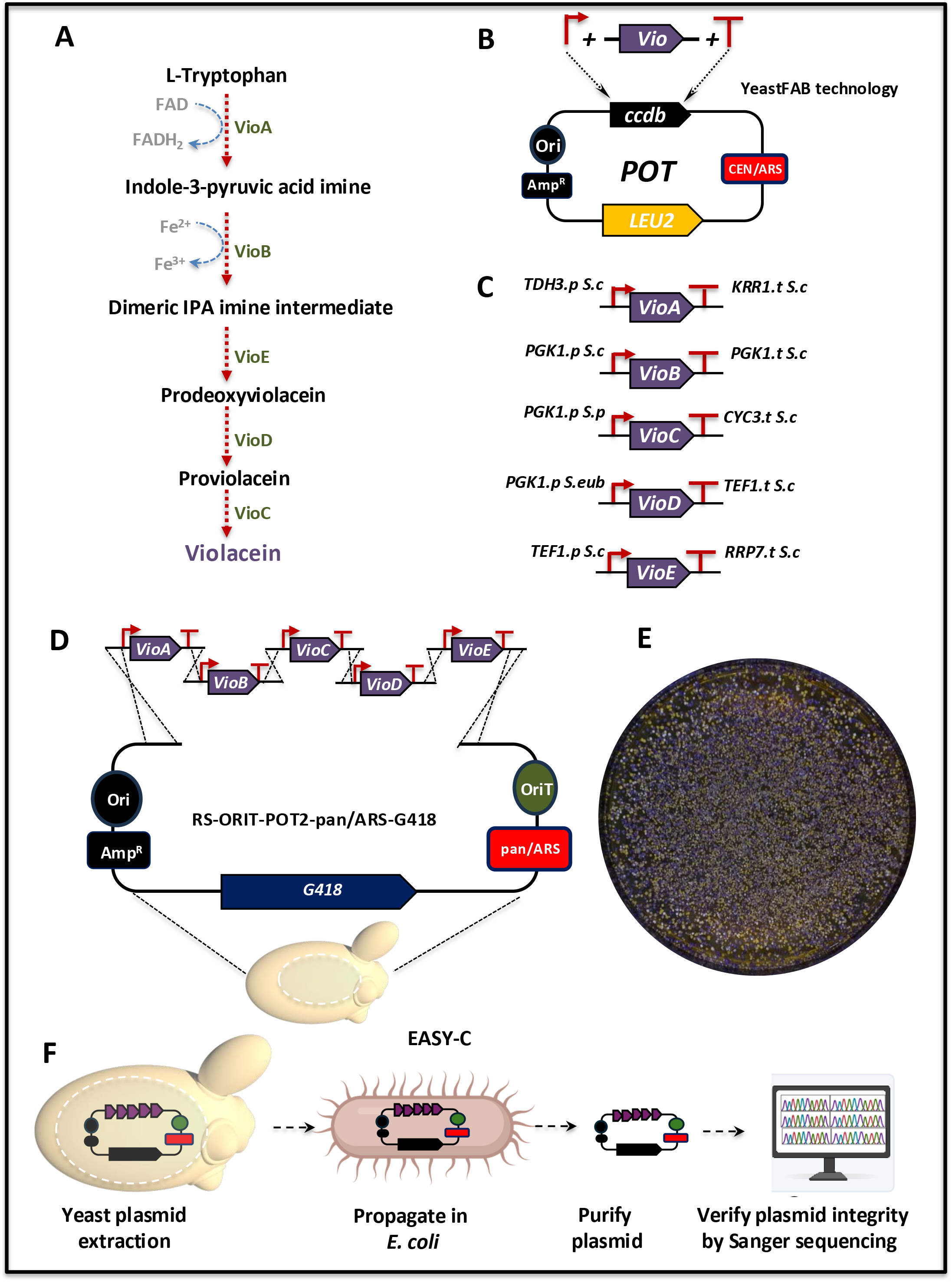
Engineering and validation of a synthetic violacein reporter for rapid visual identification of BRIDGE transformants. **(A)** Schematic of the *Chromobacterium violaceum* violacein biosynthetic pathway. L-Tryptophan is sequentially converted into violacein by the enzymes *VioA, VioB, VioE, VioD* and *VioC*, resulting in production of the characteristic purple pigment. **(B)** Construction of individual transcriptional units (promoter–ORF–terminator) using the YeastFab modular cloning system. Regulatory elements were assembled by Type IIS Golden Gate cloning, and codon-optimised *vioA–E* coding sequences were cloned to generate individual transcriptional units. **(C)** Combinatorial design of the synthetic violacein pathway. Individual transcriptional units were constructed using constitutive promoters and terminators derived from *Saccharomyces cerevisiae, Saccharomyces paradoxus* and *Saccharomyces eubayanus*. Each transcriptional unit was assembled individually into a POT vector and sequence-verified before pathway assembly. **(D)** Single-step *in vivo* assembly of the complete five-gene violacein pathway in *S. cerevisiae*. The linearised YeastFab-compatible BRIDGE broad-host delivery vector carrying Pan/ARS and oriT, together with the five transcriptional units, was assembled by homologous recombination using 60-bp overlapping regions. Positive transformants were initially identified by the appearance of purple colonies on YPD supplemented with G418 and subsequently confirmed by six junction PCR assays. **(E)** Representative *S. cerevisiae* transformants expressing the synthetic violacein pathway following selection on YPD supplemented with G418. Purple colony formation provided a rapid visual marker for identification of correctly assembled BRIDGE reporter plasmids. **(F)** EASY-C–enabled Design–Build–Recover workflow for the BRIDGE synthetic violacein reporter. Correctly assembled reporter plasmids were recovered directly from yeast using the EASY-C platform, propagated in *Escherichia coli*, purified by plasmid miniprep, and verified by full-plasmid Sanger sequencing before construction of BRIDGE donor strains for subsequent interkingdom DNA transfer.

### BRIDGE-mediated delivery and expression of the violacein pathway across the Saccharomyces genus

To evaluate the capacity of BRIDGE to transfer pathway-scale synthetic DNA, we constructed an *E. coli* donor strain carrying both the pSC5 plasmid and a BRIDGE vector harbouring the complete five-gene *vioA–E* violacein biosynthetic pathway (Materials and Methods, Section 2.6). The pathway was engineered using combinatorial constitutive regulatory elements derived from *S. cerevisiae, S. paradoxus* and *S. eubayanus* to support expression across the *Saccharomyces* genus. The resulting donor strain was used for BRIDGE-mediated interkingdom DNA transfer into six phylogenetically diverse species: *S. cerevisiae, S. paradoxus, S. mikatae, S. kudriavzevii, S. jurei* and *S. eubayanus* **(Fig. 6A, B)**. Following co-cultivation, recipient cells were selected on species-specific G418-containing medium and incubated at 30°C for 4–5 days. BRIDGE successfully delivered the complete violacein pathway to all six *Saccharomyces* species, as confirmed by colony PCR of five independent transformants from each species. Transformants were reproducibly recovered across three independent biological replicates, with the mean number of recovered transformants for each species shown in **Fig. 6E**. All six species functionally expressed the transferred pathway, producing the characteristic purple violacein phenotype, although pigmentation intensity varied between species **(Fig. 6C)**. Spot assays comparing parental strains with the corresponding BRIDGE transformants further confirmed this phenotype, with reporter-containing strains developing purple pigmentation while parental strains remained white **(Fig. 6D)**. Importantly, violacein production also provided a direct visual readout of authentic BRIDGE-mediated transfer: PCR-confirmed transformants developed purple pigmentation, whereas false-positive colonies remained white. The synthetic violacein pathway therefore provides a simple visual reporter for distinguishing authentic BRIDGE transformants while simultaneously reporting functional expression of pathway-scale synthetic DNA.

**Figure 6.**
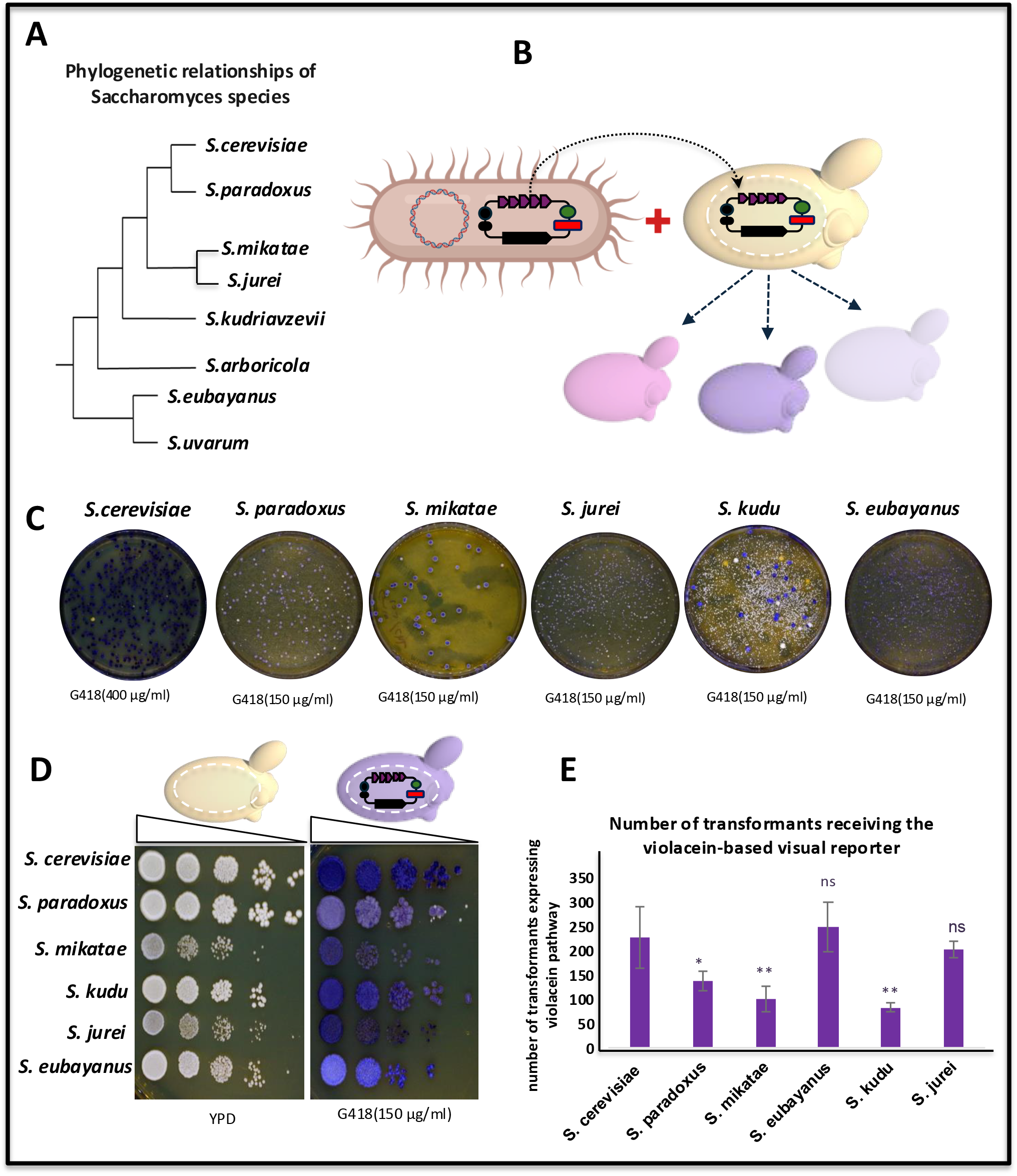
A universal violacein-based visual reporter enables rapid identification of BRIDGE transformants across *Saccharomyces* species. **(A)** Phylogenetic relationships among the six *Saccharomyces* species evaluated in this study (Montrocher, Verner et al. 1998). **(B)** Schematic of BRIDGE-mediated interkingdom DNA transfer. An *Escherichia coli* donor strain carrying the super conjugative helper plasmid and the BRIDGE synthetic violacein reporter was co-cultured with recipient *Saccharomyces* species for 24 h. Following recovery, recipient cells were selected on YPD supplemented with G418 to isolate BRIDGE transformants. **(C)** Representative BRIDGE transformants expressing the synthetic violacein reporter across six *Saccharomyces* species. Purple pigmentation varied in intensity between species but enabled rapid visual identification of positive transformants. Three independent biological replicates were performed for each species. Five independent transformants from each species were subsequently confirmed by colony PCR. **(D)** Representative spot assay comparing parental strains lacking the reporter with BRIDGE transformants carrying the synthetic violacein reporter. **(E)** Quantification of BRIDGE-mediated horizontal DNA transfer of the 16-kb synthetic violacein reporter across six *Saccharomyces* species. Bars represent the mean number of BRIDGE transformants recovered following delivery of the synthetic *vioA–E* reporter from three independent biological replicates. Error bars indicate the standard deviation (SD). Significance is indicated as follows: ns, not significant; *, *P* < 0.05; **, *P* < 0.01.

### 3.6 BRIDGE-mediated delivery and host-dependent expression of a five-gene pathway

To investigate whether BRIDGE could deliver and functionally express a complete synthetic metabolic pathway across phylogenetically diverse yeasts, an *E. coli* donor strain carrying both the pSC5 plasmid and the BRIDGE vector harbouring the violacein pathway (*vioA–E*) was used for interkingdom DNA transfer into representatives of ten yeast genera, including *S. cerevisiae, M. bulderi, S. batistae, S. bombicola, K. naganishii, K. aerobia, Y. lipolytica, Z. bailii, L. thermotolerans, K. aestuarii, P. manshurica* and *K. phaffii* (Materials and Methods, Section 2.9). Following co-cultivation, recipient cells were selected on species-specific G418-containing medium to recover transformants carrying the BRIDGE expression vector. BRIDGE-mediated delivery of the synthetic violacein reporter was evaluated using three independent biological replicates, and the mean number of recovered BRIDGE transformants is shown in **(Fig. 7A)**. Successful transformants were recovered from representatives of all ten yeast genera, demonstrating reproducible interkingdom delivery of the complete synthetic five-gene pathway across a broad phylogenetic range. However, reporter expression was host dependent. A distinguishable purple phenotype developed in *S. cerevisiae, K. aerobia* and *K. aestuarii*, whereas transformants recovered from the remaining genera exhibited little or no visible pigmentation under the conditions tested. Furthermore, pigmentation developed more rapidly and reached greater intensity in *S. cerevisiae* than in *K. aerobia* or *K. aestuarii*. To determine whether the absence of visible pigmentation resulted from unsuccessful DNA transfer or differences in pathway expression, five independent transformants from each genus were analysed by colony PCR **(Fig. 7B)**. PCR confirmed successful delivery of the BRIDGE vector carrying the complete violacein pathway in all recipient genera, irrespective of visible reporter expression. To further evaluate reporter performance, confirmed transformants were compared with their corresponding parental strains using spot assays on both rich and synthetic defined media **(Fig. 7C, D)**. Reporter-positive *S. cerevisiae* transformants displayed strong purple pigmentation, whereas *K. aerobia* developed visible pigmentation only after prolonged incubation on rich medium. Supplementation of synthetic defined medium with threefold additional L-tryptophan enhanced pigmentation in *S. cerevisiae* and accelerated colour development in *K. aerobia*, with a modest increase also observed in *K. aestuarii*. Importantly, the *vioA–E* genes were codon-optimised for *S. cerevisiae* rather than individually for each recipient species, and the pathway used regulatory elements derived from *Saccharomyces* species. Differences in codon usage and regulatory-element compatibility may therefore have contributed to the variable pigmentation observed across hosts. Collectively, these results confirm BRIDGE-mediated delivery of pathway-scale synthetic DNA across ten phylogenetically diverse yeast genera, while indicating that functional expression of the transferred pathway remains host-dependent.

**Figure 7.**
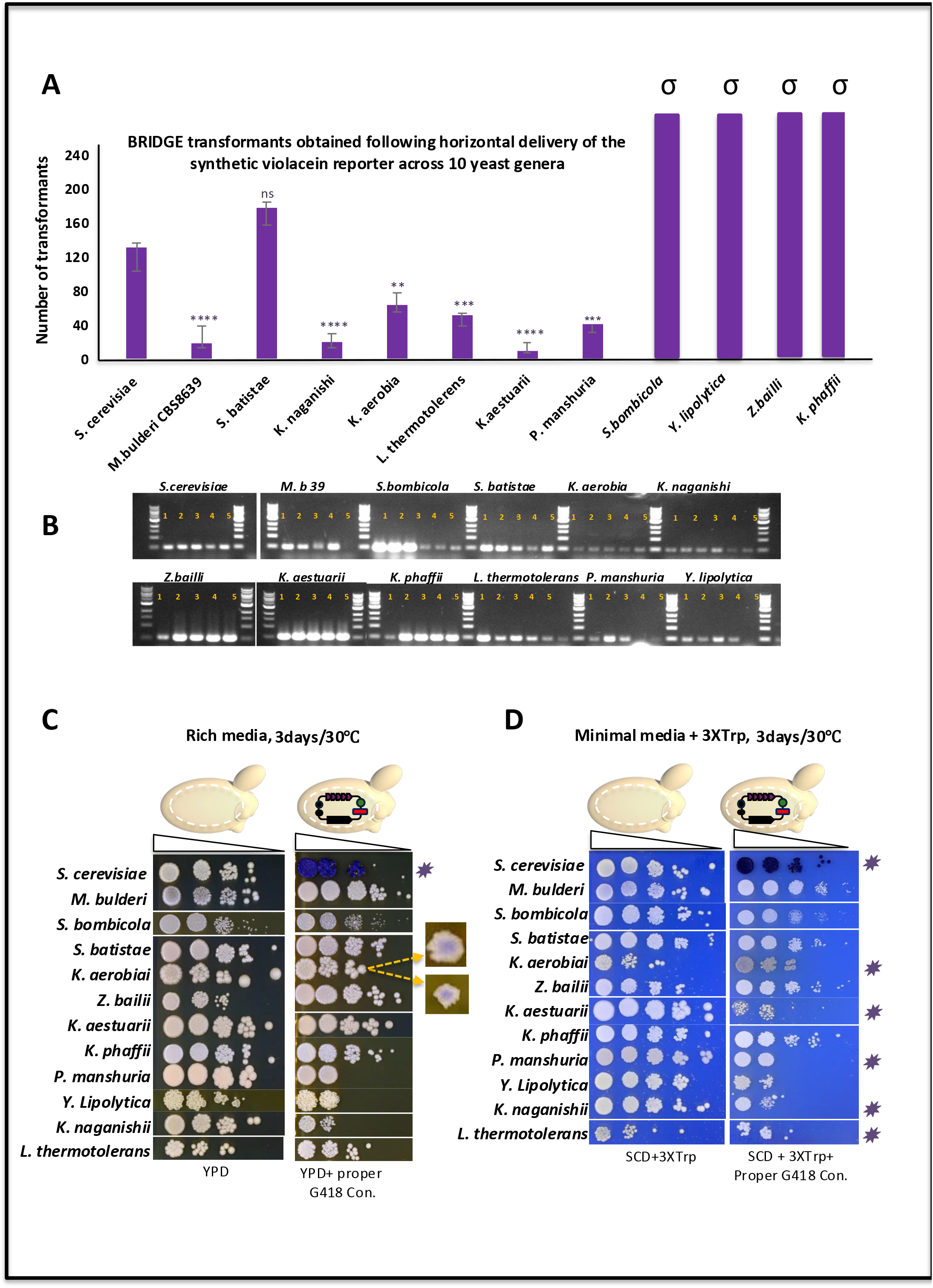
Host-dependent expression of a synthetic violacein reporter following BRIDGE-mediated delivery of a complete five-gene pathway across ten yeast genera. **(A)** Quantification of BRIDGE transformants recovered following horizontal delivery of the 16-kb synthetic violacein reporter across 12 yeast species representing 10 genera. Bars represent the mean number of BRIDGE transformants recovered from three independent biological replicates; error bars indicate the standard deviation (SD). Significance is indicated as follows: ns, not significant; **, *P* < 0.01; ***, *P* < 0.001; ****, *P* < 0.0001. Species yielding >1,000 transformants (*S. bombicola, Y. lipolytica, Z. bailii* and *K. phaffii*) formed confluent lawns rather than discrete colonies and were therefore excluded from statistical analysis. These samples are indicated by the symbol (σ) above the corresponding bars. **(B)** Colony PCR validation of five independent BRIDGE transformants from each genus following delivery of the BRIDGE broad-host delivery vector carrying the complete *vioA–E* biosynthetic pathway. **(C)** Representative spot assay on rich medium (YPD supplemented with G418) comparing parental strains lacking the BRIDGE synthetic violacein reporter with confirmed BRIDGE transformants carrying the complete *vioA–E* pathway. Reporter-positive *S. cerevisiae* transformants displayed strong purple pigmentation. In *Kazachstania aerobia*, pigmentation developed only after prolonged incubation and was initially restricted to the centre of the colony. **(D)** Representative spot assay on synthetic minimal medium supplemented with G418 and threefold additional L-tryptophan comparing parental strains and confirmed BRIDGE transformants. Supplementation with additional L-tryptophan enhanced violacein pigmentation, resulting in darker purple to nearly black colonies in *S. cerevisiae*. In *K. aerobia*, pigmentation developed more rapidly on minimal medium, producing a distinct purple-grey phenotype compared with growth on rich medium. A slight increase in pigmentation was also observed in *Kluyveromyces aestuarii*. An asterisk **(⍰)** indicates strains in which purple pigmentation or a detectable colour change was observed relative to the corresponding parental strain lacking the BRIDGE reporter under the same growth conditions.

## 3 Discussion

Horizontal gene transfer by bacterial conjugation is a major mechanism of genetic exchange and microbial evolution, facilitating the dissemination of antibiotic resistance, metabolic pathways and other adaptive traits (Frost, Leplae et al. 2005, Thomas and Nielsen 2005, Smillie, Garcillan-Barcia et al. 2010). Conjugation-mediated and related bacterial DNA-transfer systems have subsequently been applied across diverse biological systems, highlighting their potential for genetic engineering across biological boundaries. *Agrobacterium tumefaciens*-mediated transformation (ATMT), for example, is widely used for DNA delivery into plants and fungi but typically requires acetosyringone induction and co-cultivation under defined conditions (Hooykaas, van Heusden et al. 2018, Roushan, Shao et al. 2022).

In *S. cerevisiae*, a reported ATMT protocol required membrane-based co-cultivation for 6–7 days (Roushan, Shao et al. 2022). In contrast, BRIDGE requires neither acetosyringone induction nor membrane-based co-cultivation, with donor–recipient mating completed in approximately 24 h under the conditions used here. This provides a relatively simple route for exploring conjugation-mediated DNA delivery in non-conventional yeasts for which established transformation methods remain limited.

BRIDGE (Bacteria-to-Yeast Rapid Interkingdom DNA Gene Exchange) extends this concept by separating the conjugation machinery from the transferable genetic cargo. Rather than transferring the ~60-kb superconjugative helper pSC5 plasmid into recipient cells, pSC5 remains in the bacterial donor and mobilises a compact ~6-kb BRIDGE vector. The vector contains an origin of transfer (oriT) for conjugative mobilisation and a pan-fungal autonomously replicating sequence (PanARS) to support plasmid maintenance across diverse yeasts (Jayaprakash, Barroso et al. 2023). This separation reduces the size of the core transferable plasmid and, importantly, allows the genetic cargo to be changed independently of the conjugation machinery. Using this strategy, successful DNA delivery was extended beyond the previously reported fungal host range to ten phylogenetically diverse yeast genera, supporting the use of a common delivery architecture across diverse recipient species.

A second feature of BRIDGE is the modular design of the transferable vector. Rather than functioning as a fixed expression construct, the BRIDGE vector was designed as a YeastFab-compatible destination vector in which standardised promoters, ORFs and terminators can be assembled using Type IIS Golden Gate cloning (Guo, Dong et al. 2015, Garcia-Ruiz, Auxillos et al. 2018). Transcriptional units can subsequently be combined with the BRIDGE backbone through *in vivo* homologous recombination in *S. cerevisiae*, providing a practical approach for assembling larger multigene constructs. Using this strategy, the complete five-gene violacein pathway was designed, assembled, recovered and sequence-verified in less than one week. EASY-C recovery directly from yeast (Swidah, Monti et al. 2026) further enables assembled plasmids to be rapidly recovered and verified before transfer. These complementary steps form a Design–Build–Recover–Deliver (DBRD) workflow linking modular DNA construction with subsequent interkingdom delivery.

The ~10-kb five-gene violacein cargo provided a stringent test of whether BRIDGE could mobilise pathway-scale synthetic DNA rather than only individual genes or simple reporter constructs. Within the *Saccharomyces* genus, the pathway produced visible pigmentation across all six species examined, indicating that regulatory elements derived from *S. cerevisiae, S. paradoxus* and *S. eubayanus* retained substantial functionality within closely related species. Across the broader host panel, however, visible pigmentation was less consistent despite successful DNA delivery being confirmed by colony PCR. These results highlight an important distinction between DNA delivery and functional expression: successful transfer of a synthetic construct does not necessarily ensure efficient expression of its encoded pathway in a new host.

Several factors may contribute to this host-dependent expression. The *vioA–E* coding sequences were codon-optimised for *S. cerevisiae* rather than individually for each recipient species, while the regulatory elements were derived from *Saccharomyces*. Differences in codon usage, promoter and terminator compatibility, transcriptional regulation and cellular physiology may therefore influence pathway output in more phylogenetically distant yeasts (Wagner and Alper 2016, Tang, Wu et al. 2020). The increased pigmentation observed following L-tryptophan supplementation in some hosts further indicates that pathway output can also depend on metabolic context and substrate availability. These factors should therefore be considered separately from DNA-transfer efficiency when evaluating BRIDGE in new species.

Although violacein was valuable for demonstrating delivery of complex multigene cargo, it may not be the optimal primary reporter for establishing BRIDGE in an unexplored non-conventional host. Visible pigment formation requires coordinated expression and activity of five heterologous genes, making reporter output sensitive to several host-dependent variables. A simpler single-gene reporter such as GFP could provide a more direct initial assessment of functional expression. Such reporters can be codon-optimised for individual recipient species and combined with host-compatible regulatory elements, enabling rapid screening of fluorescent colonies before confirmation of DNA delivery by PCR. Importantly, the modular Golden Gate/YeastFab architecture of the BRIDGE vector allows coding sequences and regulatory elements to be exchanged without redesigning the underlying delivery system, providing a route for adapting BRIDGE to individual hosts.

This flexibility is particularly relevant for non-conventional yeasts, where the lack of efficient genetic engineering methods remains a major barrier to their exploitation (Lobs, Schwartz et al. 2017). *M. bulderi*, for example, is an acid-tolerant yeast with potential for sustainable organic acid production (Balarezo-Cisneros, Hanak et al. 2025), but remains challenging to transform. In our experience, conventional transformation requires approximately tenfold more plasmid DNA than *S. cerevisiae*, yields relatively few transformants and can generate false-positive colonies following prolonged selection, increasing experimental time and molecular validation. Once the bacterial donor carrying pSC5 and the appropriate BRIDGE vector is established, the same general mating workflow can instead be applied across different recipient yeasts, with host-specific adjustment primarily focused on appropriate selection conditions. This could reduce the need to establish new transformation procedures for every species and facilitate initial genetic access to poorly characterised hosts.

The ability to transfer the same synthetic construct across multiple species enables pathways to be assembled and verified in a tractable host before evaluation in alternative yeast backgrounds, supporting metabolic engineering, natural-product discovery and host screening. Future BRIDGE vectors could also deliver CRISPR-Cas systems, recombinases and other genome-engineering tools, alongside the development of host-compatible regulatory elements for more predictable engineering of non-conventional yeasts. Collectively, BRIDGE integrates modular DNA assembly, EASY-C plasmid recovery and broad-host interkingdom DNA delivery within a unified DBRD workflow. By separating the conjugation machinery from the transferable cargo, BRIDGE enabled DNA delivery across ten phylogenetically diverse yeast genera, including a complete five-gene biosynthetic pathway. BRIDGE therefore provides a flexible framework for expanding genetic access to non-conventional yeasts and their development as microbial cell factories for synthetic biology and industrial biotechnology.

## Supporting information

Supplementary table (Table S1): Antibiotic concentrations suitable for each yeast species

## Author contributions

**RS** conceived and designed the study with the input of **RC** and **DD. RS** performed the experimental work and wrote the manuscript. **RC** provided the super-conjugation vector and advised on the generation of donor strains. All the authors contributed to the analyses and interpretation of the data and reviewed, edited, and approved the final manuscript. **DD** and **FV** acquired the funding for this work, and **RS** was further supported by the L’Oréal-UNESCO For Women in Science UK & Ireland Rising Talent grant.

## Supplementary data

All supplementary data are available in the Supplementary Information accompanying this article. Plasmid sequences and annotated vector maps will be deposited in Addgene and will be publicly available upon publication.

## Declaration of interests

The authors declare no competing interests.

## Funding

This work was supported by the Future Biomanufacturing Research Hub (FBRH), funded by the Engineering and Physical Sciences Research Council and the Biotechnology and Biological Sciences Research Council (grant number EP/S01778X/1), and by the L’Oréal-UNESCO For Women in Science UK & Ireland Rising Talent grant awarded to **RS**.

## Acknowledgements

The authors would like to thank Professor Anthony Green, Director of the Manchester Institute of Biotechnology, and Professor Chris Hardacre, Head of the School of Natural Sciences at The University of Manchester, for their support of R.S. and her research.

## Figure legends

**Supplementary Figure 1.**
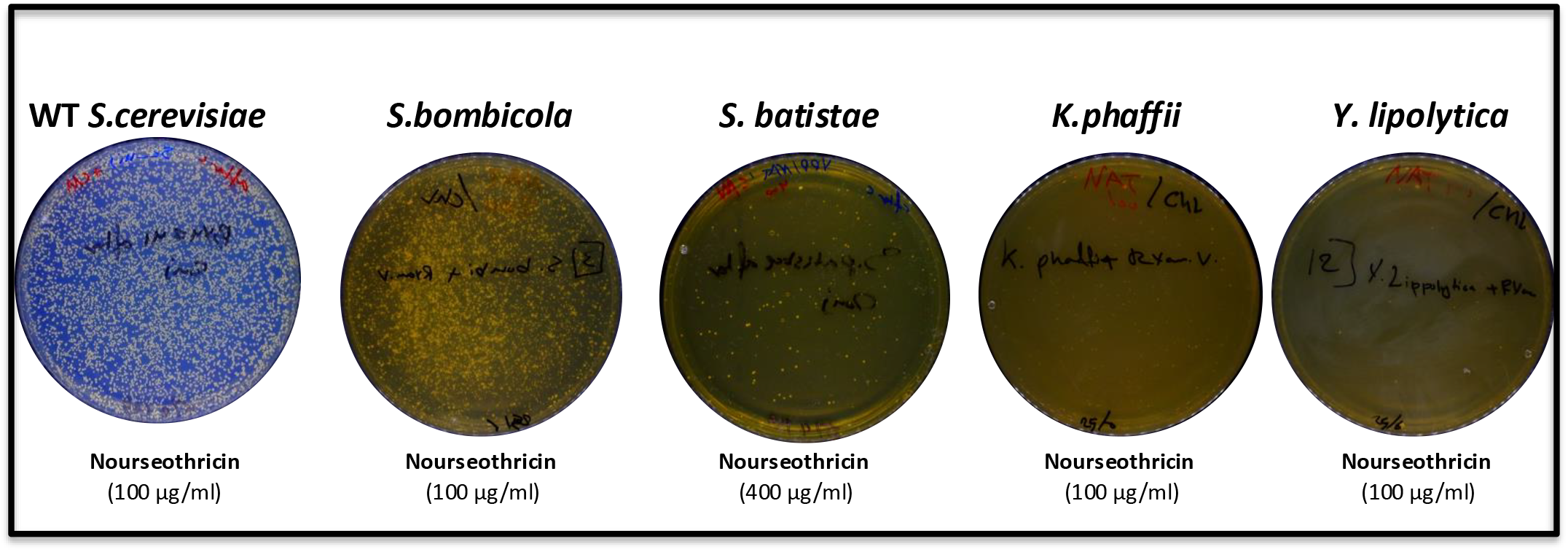
Transformants obtained following horizontal transfer of the super conjugative plasmid. Representative transformants obtained four days after horizontal transfer of the superconjugative helper plasmid into *Saccharomyces cerevisiae, Starmerella batistae, Starmerella bombicola, Yarrowia lipolytica*, and *Komagataella phaffii*. Transformants were selected using nourseothricin to select for yeast transformants and chloramphenicol to eliminate residual *E. coli* donor cells. Representative transformants from each species were confirmed by colony PCR.

**Supplementary table (Table S1):**
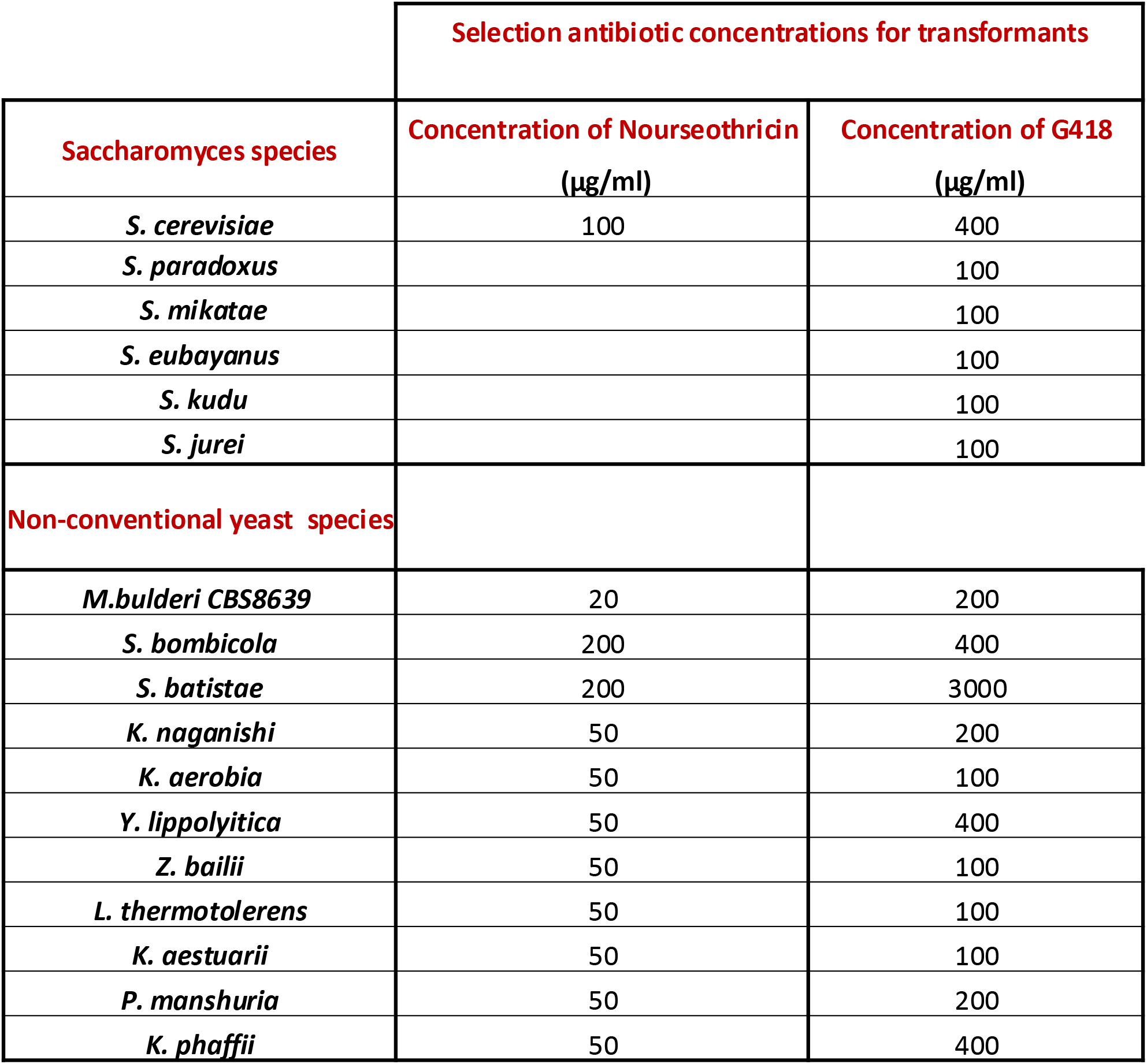
Antibiotic concentrations suitable for each yeast species.

## Notes

### Competing Interest Statement

The authors have declared no competing interest.

