## Supplementary table (Table S1): Antibiotic concentrations suitable for each yeast species for "BRIDGE: A broad-host platform for interkingdom DNA delivery enabling the genetic domestication of phylogenetically diverse yeasts"

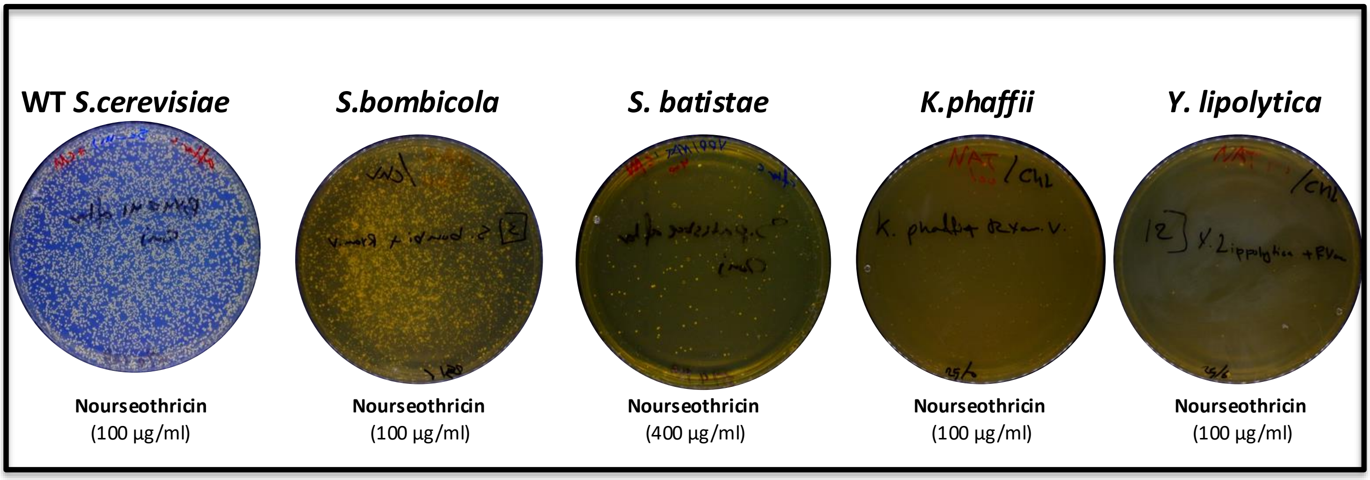


**Supplementary Figure 1. Transformants obtained following horizontal transfer of the superconjugative plasmid.**

Supplementary table (Table S1): Antibiotic concentrations suitable for each yeast species

| | Selection antibiotic concentrations for transformants | |
|---|---|---|
| **Saccharomyces species** | **Concentration of Nourseothricin (µg/ml)** | **Concentration of G418 (µg/ml)** |
| ***S. cerevisiae*** | 100 | 400 |
| ***S. paradoxus*** | | 100 |
| ***S. mikatae*** | | 100 |
| ***S. eubayanus*** | | 100 |
| ***S. kudu*** | | 100 |
| ***S. jurei*** | | 100 |
| **Non-conventional yeast species** | | |
| ***M.bulderi CBS8639*** | 20 | 200 |
| ***S. bombicola*** | 200 | 400 |
| ***S. batistae*** | 200 | 3000 |
| ***K. naganishi*** | 50 | 200 |
| ***K. aerobia*** | 50 | 100 |
| ***Y. lippolyitica*** | 50 | 400 |
| ***Z. bailii*** | 50 | 100 |
| ***L. thermotolerens*** | 50 | 100 |
| ***K. aestuarii*** | 50 | 100 |
| ***P. manshuria*** | 50 | 200 |
| ***K. phaffii*** | 50 | 400 |
